# *Bifidobacterium castoris* strains isolated from wild mice show evidence of frequent host switching and diverse carbohydrate metabolism potential

**DOI:** 10.1101/2021.06.03.446870

**Authors:** Magdalena Kujawska, Aura Raulo, Laima Baltrūnaitė, Sarah CL Knowles, Lindsay J Hall

## Abstract

Members of the gut microbiota genus *Bifidobacterium* are widely distributed human and animal symbionts believed to exert beneficial effects on their hosts. However, in-depth genomic analyses of animal-associated species and strains are somewhat lacking, particularly in wild animal populations. Here, to examine patterns of host specificity and carbohydrate metabolism capacity, we sequenced whole genomes of *Bifidobacterium* isolated from wild-caught small mammals from two European countries (UK and Lithuania). Members of *B. castoris, B. animalis* and *B. pseudolongum* were detected in wild mice (*Apodemus sylvaticus, A. agrarius* and *A. flavicollis*), but not voles or shrews. *B. castoris* constituted the most commonly recovered *Bifidobacterium* (78% of all isolates), with the majority of strains only detected in a single population, although populations frequently harboured multiple co-circulating strains. Phylogenetic analysis revealed that the mouse-associated *B. castoris* clades were not specific to a particular location or host species, and their distribution across the host phylogeny was consistent with regular host shifts rather than host-microbe codiversification. Functional analysis suggested that mouse-derived *B. castoris* strains encoded an extensive arsenal of carbohydrate-active enzymes, including putative novel glycosyl hydrolases such as chitosanases that may act on chitin-derived substrates such as mushrooms or insects, along with genes encoding putative exopolysaccharides, some of which may have been acquired via horizontal gene transfer. Overall, these results provide a rare genome-level analysis of host specificity and genomic capacity among important gut symbionts of wild animals, and reveal that *Bifidobacterium* has a labile relationship with its host over evolutionary time scale.

## Introduction

Species and strains belonging to the bacterial genus *Bifidobacterium* are prominent members of the gut microbiota in many animals, and are universally distributed among animals exhibiting parental care, including humans and non-human mammals, birds and social insects (*1*). *Bifidobacterium* species that colonise the human gut, especially those associated with early life stages, have received much attention in recent years due to their ability to confer health benefits on their host, including supporting development of the wider gut microbial ecosystem through the production of short chain fatty acids, colonisation resistance against pathogens and immune modulation. These beneficial properties have been linked to their carbohydrate metabolism and exopolysaccharide (EPS) biosynthesis capabilities (*2, 3*).

At the time of the writing (January 2021), over 80 *Bifidobacterium* species and subspecies have been identified in multiple animal hosts, with around 2,400 genome assemblies available via the NCBI Genome database (*4*). However, the majority of these sequences come from human-associated species, namely; *Bifidobacterium longum, Bifidobacterium breve, Bifidobacterium bifidum*, and *Bifidobacterium pseudocatenulatum*. Consequently, strains belonging to these species, and single strains that represent type strains (some of which were originally isolated from captive animals), comprise the majority of genomes available, and are thus the best studied with respect to comparative genomics (*1, 5-10*). Therefore, our current knowledge of the diversity and evolution of *Bifidobacterium* within the mammalian gut remains limited.

Reports of congruent phylogenies between mammals and gut microbial taxa have suggested that animal hosts and their symbionts may diversify and speciate together (*11-14*). Codiversification patterns have been linked to convergent acquisition of function by different bacterial phylogenetic clades, with horizontal gene transfer and gene loss proposed as potential mechanisms involved in the process (*15, 16*). Furthermore, increased host-specificity has been suggested to be linked to reduced transmission capacity due to anaerobic and non-spore forming bacterial lifestyles (*17, 18*). As gut microbes can affect host phenotype in a variety of ways, for example by modulating host energy acquisition and immune development (*19, 20*), understanding and documenting the distribution and diversification patterns of key bacterial species within hosts constitutes an important step towards understanding host-microbe interactions over evolutionary time (*21*).

In *Bifidobacteriaceae* specifically, phylogenetic congruence between four great ape species and their *Bifidobacteriaceae* gyrB lineages suggested a significant degree of codiversification, albeit with some host switching (*11*). Furthermore, the analysis of cophylogenetic relationships between 24 primate hosts and 23 primate-associated bifidobacterial species revealed the existence of phylogenetic congruence between *Bifidobacterium* typically associated with human hosts (*B. adolescentis, B. bifidum, B. breve, B. catenulatum, B. dentium, B. longum* spp. and *B. pseudocatenulatum*) and the members of Hominidae family (*Gorilla gorilla, Homo sapiens* and *Pan troglodytes*) (*22*). In contrast, insights into genotypic and phenotypic properties of animal-associated *Bifidobacterium animalis* and *Bifidobacterium pseudolongum* revealed that strains belonging to these species were found in animal hosts spanning both mammals and birds, and thus appear to be generalist at the species level, rather than having host-specific niches (*23, 24*). The widespread distribution of different *Bifidobacterium* species at higher taxonomic ranks of the mammalian tree of life (family and order) has also been suggested based on the analysis of the short internally transcribed spacer (ITS) rRNA sequences (*25*). However, since robust strain-level information on the majority of animal-associated *Bifidobacterium* species is lacking, it remains unclear to what extent the proposed generalism holds true at higher taxonomic resolution, and whether highly resolved strains are in fact host specific. This is particularly important when exploring potential adaptation of bacterial symbionts to their hosts in wild populations (rather than model organisms or captive animals), because the adaptive relationship is dependent on the environmental niche, to which the host (and potentially gut microbe) have adapted over evolutionary time scale.

It remains unclear which processes govern the evolution of mammalian gut microbial symbionts (*26, 27*). Recently, Groussin et al. (*28*) proposed that reciprocal and specific functional dependencies between mammalian hosts and single bacterial clades are not strong enough for coevolution to occur and drive cospeciation events. Instead, the authors proposed that allopatric speciation, which implies geographic isolation of the host species and subsequent limited symbiont dispersal and diversification, may lead to host-symbiont cospeciation and phylogenetic congruence patterns. According to this model, host adaptation to new conditions following an allopatric event can result in an altered intestinal environment, to which symbiotic bacteria can then quickly adapt (e.g., different glycan composition of plant-derived substrates in herbivore hosts) (*28*). While studies using methods of broad taxonomic resolution (e.g. down to genus- or species-level) provide some information on evolutionary relationships between hosts and their gut microbes, studies involving high (strain-level, or genomic) taxonomic resolution in natural systems are lacking, yet may reveal cryptic diversity and patterns of host specificity and host-microbe evolution not previously appreciated, and shed light on the functional role that particular bacterial symbionts play in the wider gut microbiota of wild animals.

Wild small mammals provide an excellent study system in which to explore host-gut microbe relationships in more detail, as they are geographically widespread, diverse, easily trapped and possess a rich gut microbiota. To bridge the current knowledge gap on the distribution of *Bifidobacterium* in wild animals, we chose wild rodents as a target host group in which to profile patterns of *Bifidobacterium* diversity, and determine strain-level and potential functional adaptation to different host species and geographical regions. To this end, we surveyed wild mice, voles and shrews from multiple populations in two geographically distinct parts of Europe, performed *Bifidobacterium* isolations, and subsequently investigated a collection of derived *Bifidobacterium* genomes. Phylogenomic and functional genomic analysis indicated enrichment for *B. castoris* and particular carbohydrate metabolism and host modulatory properties.

## Results

Between December 2015 and December 2018, we collected and processed 220 faecal samples from 9 species of small mammals (mice, voles and shrews) caught at 14 sites across two European countries – Lithuania and the UK. In the UK, samples were collected from two mouse species (*Apodemus sylvaticus* and *Apodemus flavicollis*) in both Wytham Woods (Oxfordshire, n=78) and Silwood Park (Berkshire, n=14). In Lithuania, 54 samples from mice (*Apodemus agrarius, A. flavicollis*), 61 samples from voles (*Microtus agrestis, Microtus arvalis, Microtus oeconomus, Myodes glareolus*) and 13 samples from shrews (*Sorex araneus, Sorex minutus, Neomys fodiens*) were obtained across 12 trapping sites (**Supplementary Table S1**).

Our isolation efforts resulted in the recovery of 51 *Bifidobacterium* isolates from a total of 32 individuals belonging to three wild mouse species – *A. flavicollis, A. sylvaticus*, and *A. agrarius. Bifidobacterium* was isolated from 21.9% of mouse samples screened, including 27.0% *A. flavicollis* samples, 26.3% *A. sylvaticus* samples and 6.9% *A. agrarius* samples, respectively. We were not successful in recovering *Bifidobacterium* from voles or shrews. The probability of isolating *Bifidobacterium* varied strongly across host families (Pearson’s *X*^2^ (df=2) = 18.98, *P* < 0.001). Whole genome sequencing of recovered isolates yielded a mean of 265-fold coverage for samples sequenced on HiSeq (minimum 172-fold, maximum 300-fold) and 225-fold for samples sequenced on MiSeq (minimum 130-fold, maximum 325-fold). One sequence did not assemble correctly and was removed from further analysis. Based on the literature, we defined sequences exhibiting the average nucleotide identity (ANI) value > 99.9% as identical (*29*). Using this threshold, we excluded further 17 duplicate genomes representing identical isolates from the same individuals sequenced multiple times. This resulted in the final dataset comprised of 33 *Bifidobacterium* genomes representing isolates recovered from 31 individual hosts. In total, we identified 26 isolates as *B. castoris*, 4 isolates as *B. animalis* and a further 3 isolates as *B. pseudolongum* (**Supplementary Table S1 and Supplementary Table S3**).

The assembled draft genome sizes for mouse-associated *B. castoris* ranged from 2.27 Mb to 2.39 MB, possessing an average G+C% content of 65.53% and a number of contigs ranging from 9 to 56. The number of predicted ORF in each genome ranged from 1,832 to 1,980. Genome size and gene number was therefore lower in comparison to the only current genome sequence available (January, 2021) for the type strain *B. castoris* 2020B^T^ (GenBank accession: GCA_003952025.1), isolated from a captive beaver (*Castor fiber*) in Italy, whose genome size was 2.50 Mb, with 2,053 ORFs and an average G+C% content of 65.41% (*30*). The sizes of draft genomes for *B. animalis* and *B. pseudolongum* ranged from 2.15 Mb to 2.19 Mb (1,808 to 1,849 ORFs) and 2.03 Mb to 2.06 Mb (1,705 to 1,725 ORFs), respectively. *B. animalis* strains had an average G+C% content of 60.00%, while this value was at 63.34% for *B. pseudolongum*. These findings are in line with previous reports for members of these species isolated from rodents (*23, 24*).

In terms of *Bifidobacterium* distribution across host species, *A. sylvaticus* (n= 19, UK) was found to harbour *B. castoris* and *B. animalis. A. flavicollis* in the UK (n=2) only harboured *B. animalis*, whereas the same host species in Lithuania (n=8) harboured *B. castoris* and *B. pseudolongum. A. agrarius* (n=2, Lithuania) was found to only harbour *B. castoris*. Based on the previously set ANI threshold (ANI > 99.9%) (*29*), we identified five *B. castoris* and two *B. animalis* strains in *A. sylvaticus*, five *B. castoris*, two *B. animalis* and three *B. pseudolongum* in *A. flavicollis*, and two *B. castoris* strains in *A. agrarius*. Overall, our isolation efforts resulted in the recovery of 12 *B. castoris*, 4 *B. animalis* and 3 *B. pseudolongum* strains. On average, we recovered one unique *Bifidobacterium* strain per individual, except for one *A. sylvaticus* individual from Wytham (X0418EBC072), who was found to harbour both *B. castoris* and *B. animalis*. However, a much larger sequencing effort per sample would be required to assess the prevalence and the abundance of multiple species and strains in individual hosts.

Interestingly, all newly sequenced isolates belonged to the previously established *B. pseudolongum* phylogenetic group (*31*). Recent taxonomic analyses of the genus *Bifidobacterium* indicated that this phylogenetic group was very diverse in terms of ecological niches represented by host species, and encompassed strains isolated from animals as diverse as chickens, geese, dogs, oxen, pigs, rabbits, hamsters and rats (*31*). Since the relatedness of organisms can effectively be predicted based on their shared gene content (*32, 33*), we constructed a maximum-likelihood phylogenetic tree using single copy core genes (n=610) to assess relationships between our isolates and representative members of the *B. pseudolongum* group (n=112), with a particular focus on strains isolated from rodents (**Figure 1 and Supplementary Table S1**). Despite the limited number of rodent-associated *Bifidobacterium* genomes available for this analysis (n=20), results indicate some clustering of strains according to host phylogeny. For example, while *B. animalis* isolates recovered from mice tend to cluster separately from those from rats, *B. pseudolongum* isolates from a porcupine and a patagonian mara cluster together. This observation is in line with previous genomic analysis of animal-associated *B. pseudolongum* isolates, which indicated that different animal hosts harbour specific clusters of members of this taxon (*24*).

**Figure 1.**
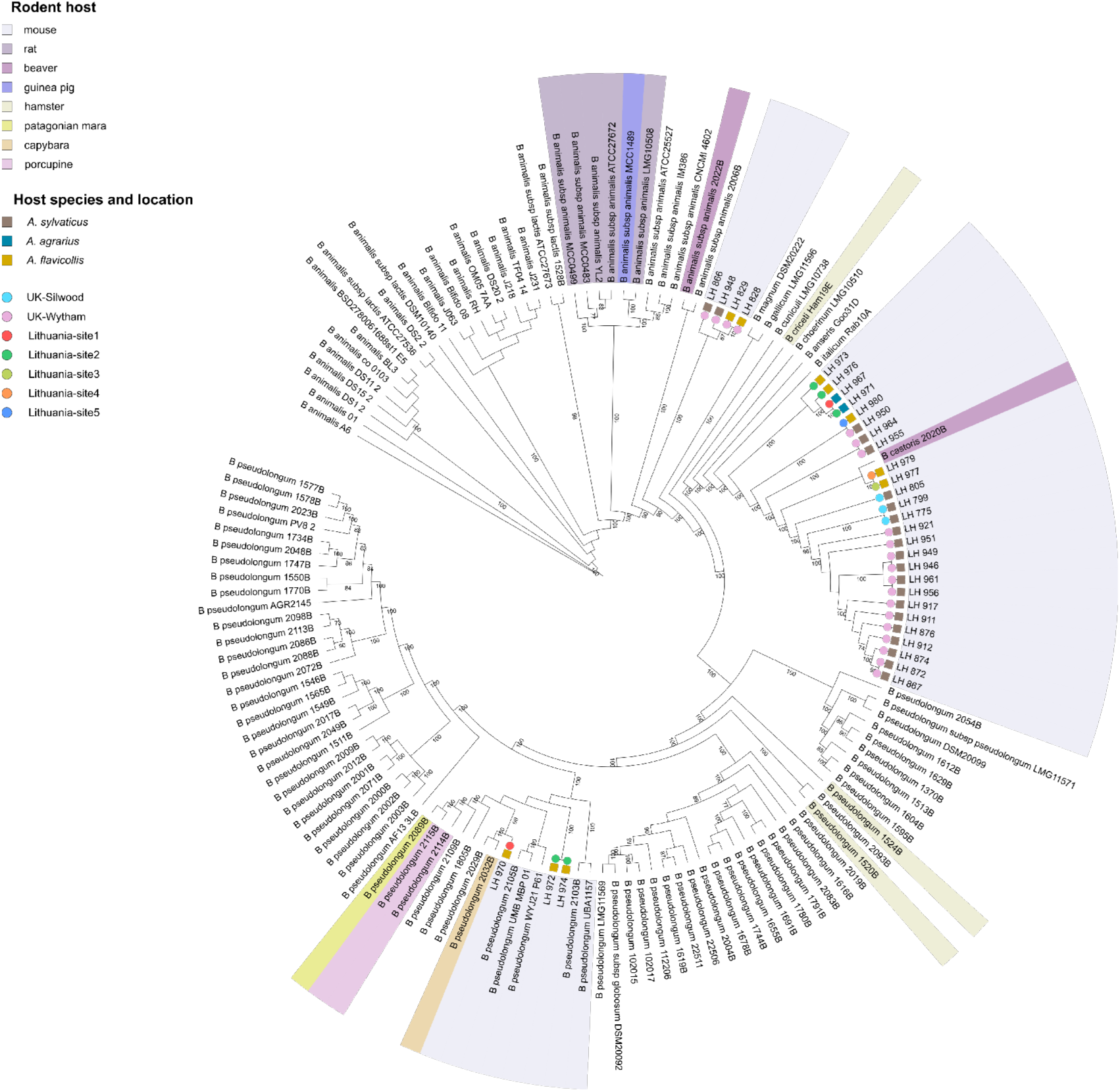
Cladogram of *B. pseudolongum* phylogenetic group, including 112 publicly available representative strains and the 33 isolates recovered in this study. Maximum likelihood phylogeny was based on protein sequences of single copy core genes (n=610), employing the ‘WAG’ general matrix model with 1000 bootstrap iterations. Bootstrap values above 70% are displayed on tree branches. Strains isolated from rodent hosts are marked with coloured background. Coloured symbols on the branches depict respective host species (square) and trapping sites (circle) for isolates recovered in this study.

Since *B. castoris* constituted 78% of all *Bifidobacterium* isolates recovered in this study, and this species is the least well-characterised, we focused further detailed genomic analyses on this species. The pangenomic analysis of these isolates alongside the type strain previously isolated from a beaver revealed a total of 2,897 gene clusters (**Figure 2**). Based on the distribution of gene clusters in the pangenome, we identified 1,412 gene clusters that constituted the core genome shared by all isolates (48.7% of all clusters), while 438 clusters (15.1% of all clusters) were unique genes (**Supplementary Table S2**). Using protein sequences for single copy core genes of the pangenome we constructed a *B. castoris* phylogeny based on maximum likelihood estimation (**Figure 2 and Figure 3a**).

**Figure 2.**
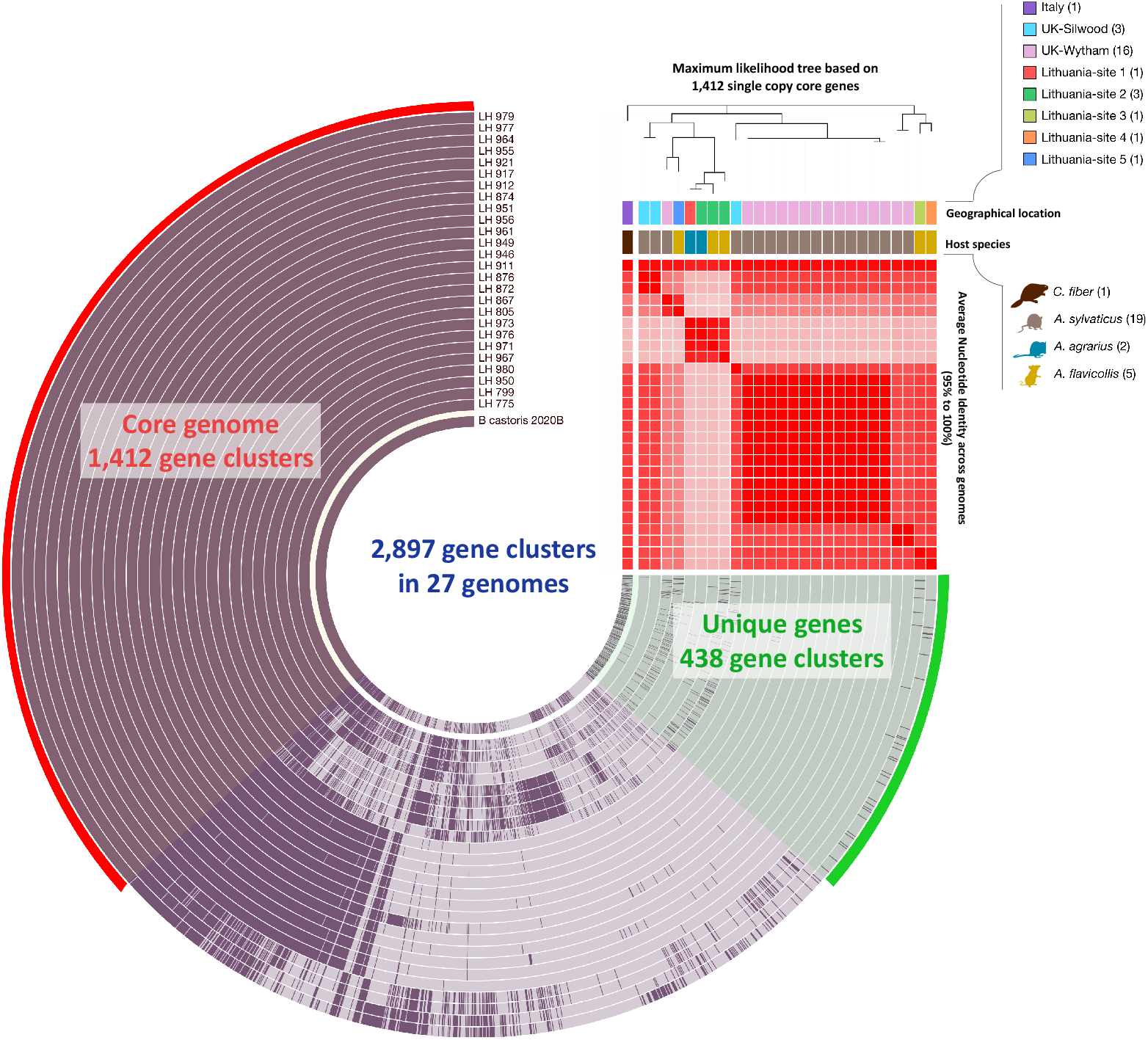
Pangenomic analysis of 27 genomes of *Bifidobacterium castoris* revealing 1,412 (48.7% of all clusters) core gene clusters, and 438 (15.1%) strain-specific unique gene clusters among 2,897 total gene clusters, along with their distribution and average nucleotide identities (ANI > 95%).

**Figure 3.**
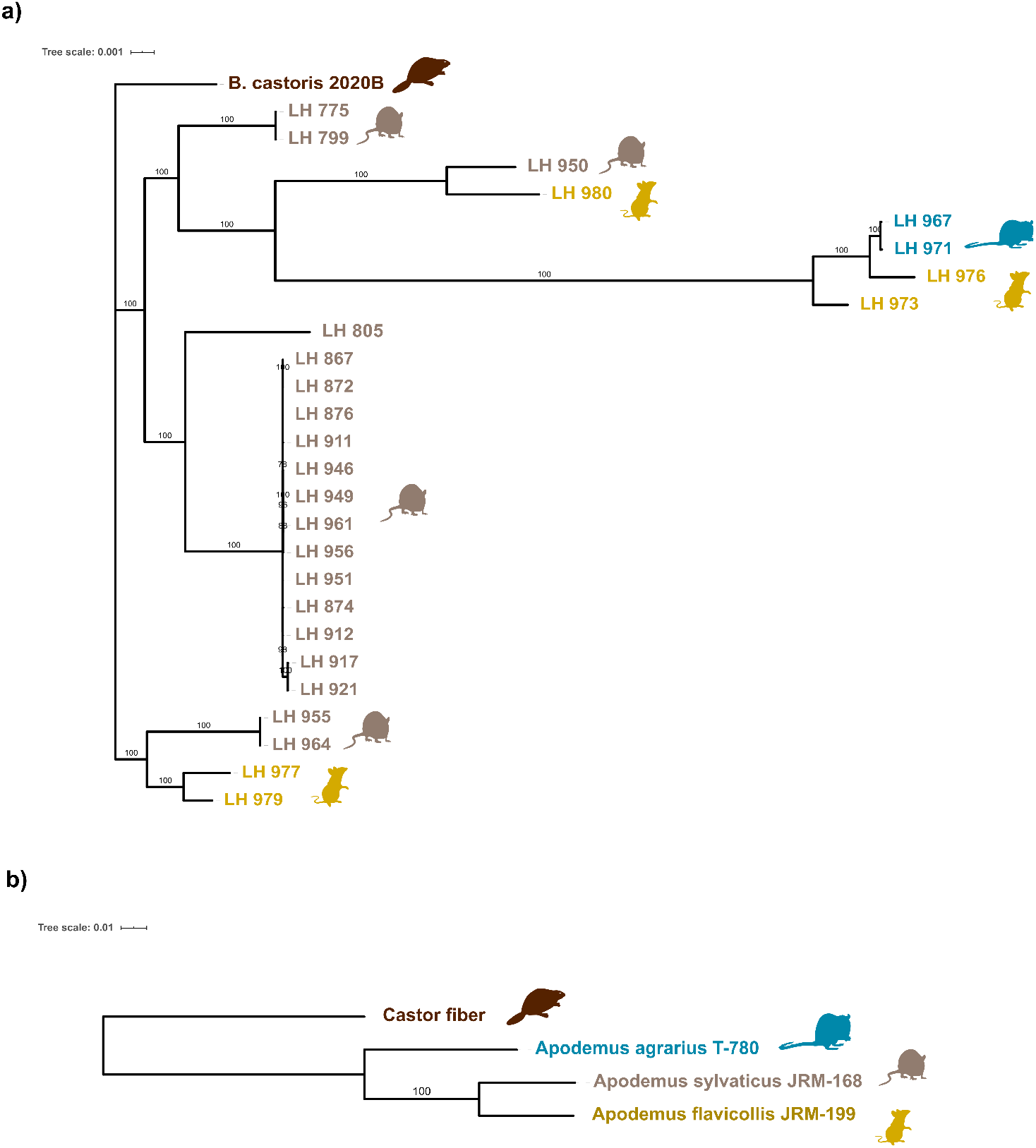
Phylogeny of 27 *Bifidobacterium castoris* isolates (a) and their rodent hosts (b). Maximum likelihood trees were constructed using single copy core genes employing ‘WAG’ model and 1000 bootstrap iterations for *B. castoris* and concatenated 12S rRNA and partial cytochrome b genes employing ‘GTR’’model with 1000 bootstrap iterations for host species. Bootstrap values above 70% are displayed on tree branches.

Examination of the *B. castoris* phylogenetic tree (**Figure 3a**), revealed that the 12 mouse-isolated strains fell into three major clusters. One cluster is more distant from the other two and contains strains from both *A. sylvaticus* and *A. flavicollis*. The second cluster seems to be *A. sylvaticus*-specific and appears to contain 2 main strains, while the third main cluster contains strains isolated from all three *Apodemus* species, including one *A. sylvaticus*-specific subclade, and another subclade that contains two clusters of strains that each colonise two different host species. Overall, these observations are not consistent with a strong pattern of cospeciation. Therefore, we sought to test for cophylogenetic signal between *Bifidobacterium* isolates and their hosts (*34*). The host tree was constructed using concatenated sequences for part of the cytochrome b (cytb) gene and the mitochondrial 12S rRNA gene (**Figure 3b**) (*35*). Cophylogeny was first tested for using the global ParaFit statistic (H0: *B. castoris* and its hosts have independent phylogenetic structure). The result (permutational *P* = 0.3586 after 9999 permutations) was not significant, failing to reject the null hypothesis, and thus providing no evidence that the phylogeny of *B. castoris* and its hosts are correlated. Further ParaFit test of associations between individual *Bifidobacterium* isolates and their respective hosts did not reveal any significant links (*P* > 0.05, **Table 1**), suggesting the absence of co-phylogenetic patterns.

With five *B. castoris* strains identified in *A. sylvaticus*, five in *A. flavicollis*, and two in *A. agrarius*, we next sought to determine how many different strains circulate in each country. Overall, a total of 5 *B. castoris* strains were detected across the two UK sites: three distinct strains in Wytham, with one strain more common than the others (detected 13 times), and two strains at Silwood Park. In Lithuania, only one site showed evidence of more than one strain circulating (two strains detected in *A. flavicollis* and one strain detected in *A. agrarius* at site2). No strains were found in more than one murine host species nor multiple locations. For the only strain that we detected fairly regularly (Strain 3 LH_867-LH_961, found 13 times in *A. sylvaticus* in Wytham), we tested whether prevalence varied by host species or geographical location. This strain’s occurrence differed significantly across host species and countries (which are confounded), as it was only detected in UK wood mice (Fisher’s exact tests for both host species and country, p=0.005). Altogether, these results suggest that *B. castoris* strains display a certain degree of species-and site-specificity.

To determine functional differences between *B. castoris* isolates, we next functionally annotated ORFs of each genome based on orthology assignment using eggNOG-mapper. This analysis resulted in the classification of an average of 82.46% genes per genome into COG categories, and reflected the saccharolytic lifestyle of *B. castoris*, with carbohydrate transport and metabolism identified as the second most abundant category (after unknown function) constituting 9.98% of functionally annotated genes. This value is slightly higher compared to previous findings for the pangenome of animal-associated *B. pseudolongum* taxon (9%) and within the range reported for other bifidobacteria (*24, 36*) (**Supplementary Table S4**). Members of *Bifidobacterium* have been shown to synthesize and digest a wide range of carbohydrates through an extensive arsenal of carbohydrate-active enzymes (CAZymes) (*37, 38*). We thus sought to investigate the genetic repertoire predicted to be involved in carbohydrate metabolism and biosynthesis in *B. castoris. In silico* analyses performed using dbCAN2 identified three classes of enzymes, namely glycosyl hydrolases (GHs), glycosyl transferases (GTs) and carbohydrate esterases (CEs), as well as enzyme-associated carbohydrate-binding modules (CBMs) (**Figure 4a and Supplementary Table S5**). On average, *B. castoris* genomes harboured 86.52±3.15 CAZymes. Previous reports on CAZyme abundances in strains isolated from different hosts and environments showed that, on average, *Bifidobacterium* isolated from rodents had less than 50 CAZyme genes in their genomes, a number comparable with strains isolated from dairy and wastewater (*39*). Interestingly, the abundance of CAZymes similar to that of *B. castoris* was reported for strains isolated from non-human primates (84±10 CAZymes) (*39*).

**Figure 4.**
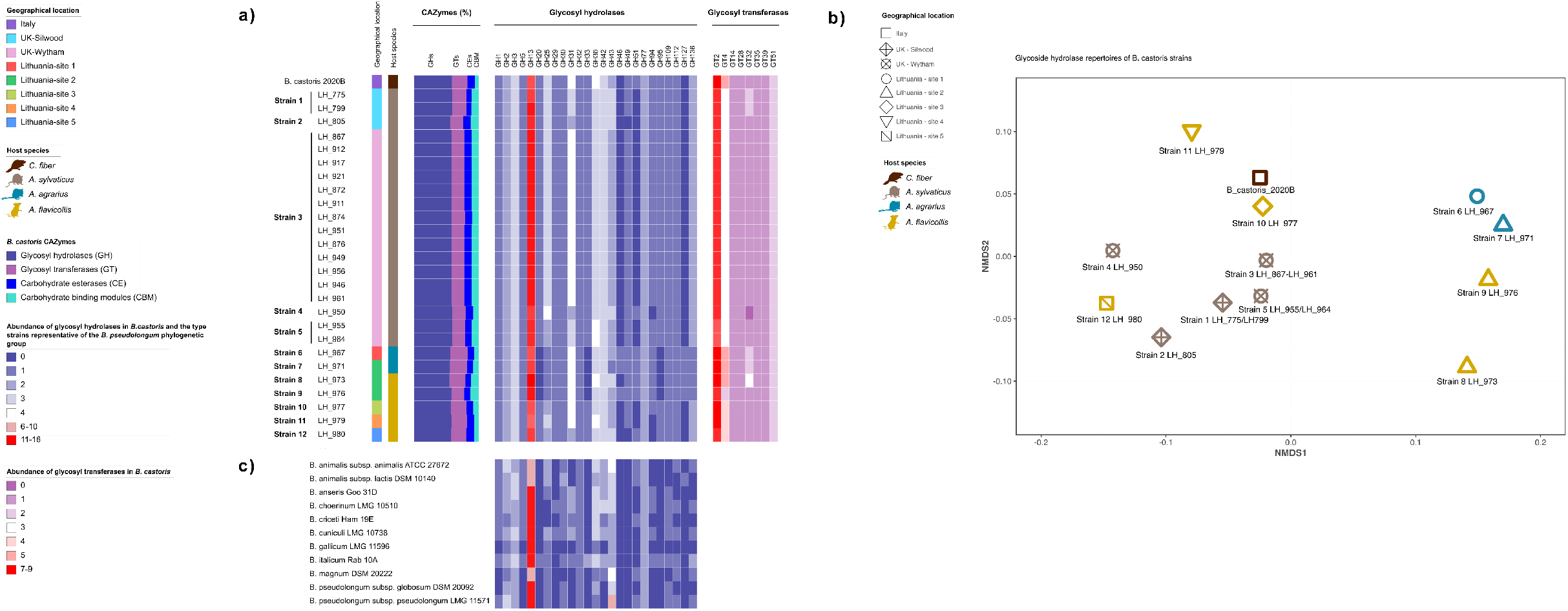
Predicted glycobiome of *B. castoris* (a), clustering of GH profiles of *B. castoris* strains using non-metric multidimensional scaling (NMDS) (b) and the abundance of glycosyl hydrolases in *Bifidobacterium* type strains representative of the *B. pseudolongum* phylogenetic group (c). (a) Coloured strips indicate geographical location and host species. The bar plot (CAZymes %) shows proportional representation of four CAZyme classes constituting the predicted *B. castoris* glycobiome. The two heatmaps present abundances of genes predicted to belong to specific glycosyl hydrolase (GH -blue) and glycosyl transferase (GT -purple) families. (b) The heat maps shows abundances of genes predicted to belong to the same glycosyl hydrolase families as those identified in *B. castoris*.

Glycosyl hydrolases are key enzymes in carbohydrate metabolism that catalyse the hydrolysis of glycosidic bonds between two or more carbohydrates or between a carbohydrate and non-carbohydrate moiety (*40*). We identified a total of 25 different GH families in *B. castoris* isolates containing an average of 49.48±1.99 GH genes per genome (2.62%±0.16 of ORFs and 57.68±0.01% of predicted glycobiome). The predominant GH family, with 14.15±1.29 GH genes per genome (mean±sd), constituting on average 29.98±1.88% of the GH repertoire, was GH13 whose members include enzymes acting on a very wide range of carbohydrates containing α-glucoside linkages, e.g. starches and related substrates, trehalose, raffinose, stachyose and melibiose (*9, 38, 41*). These results were consistent with previous studies of the type strains representative of the genus *Bifidobacterium*, as well as strains belonging to *B. pseudolongum* species, which identified this particular GH family as the most commonly detected, in particular in the genomes of strains isolated from mammals (*9, 24*). Families GH31 (a diverse group of enzymes with α-glucosidase and α-xylosidase activities) and GH36 (enzymes metabolising α-galacto-oligosaccharides present in various plants, i.e. melibiose, raffinose, stachyose) (*42, 43*) followed, with 3.63±0.68 and 3.18±0.39 GH genes per genome, respectively.

It has been well established that the GH repertoires of *Bifidobacterium* are species-and often strain specific (*9, 23, 24, 44*). Therefore, we expected that the diverging *B. castoris* strains would also harbour diverging GH profiles. Indeed, at every geographical location, the isolates identified as unique strains also displayed individual GH profiles (**Figure 4a and b, and Supplementary Table S5**). However, we noted one instance of an inconsistent prediction of the number of genes belonging to GH13 family between the two isolates identified as Strain 1 from Silwood (13 vs 15 GH genes in LH_775 and LH_799, respectively), with identical values for all other identified GH families. This result may possibly be explained by the differences in the number of contigs between the genomes of these isolates and the subsequent differences in the annotation.

The comparison between GH profiles of *B. castoris* (**Figure 4a and Supplementary Table S5)** and those of 9 *Bifidobacterium* type strains representative of the members of the *B. pseudolongum* phylogenetic group (**Figure 4c and Supplementary Table S5**) revealed that *B. castoris* species alone appears to possess GH49 family, which contains dextranases acting on dextran and pullulan (*45*). The majority of *B. castoris* strains isolated from hosts across all trapping sites appear to be lacking families GH51 and GH127, but possess several copies of genes predicted to encode enzymes belonging to GH43 family. These GH families predominantly contain α-and β-L-arabinofuranosidases that hydrolyse the glycosidic bond between L-arabinofuranoside side chains of hemicelluloses such as arabinoxylan, arabinogalactan and L-arabinan, naturally present in cereal grains (*46, 47*). The exception are 4 strains isolated from *A. agrarius* and *A. flavicollis* from site1 and site2 in Lithuania, 3 out of which appear to additionally possess predicted GH46 chitosanases acting on chitin-derived substrates (i.e. mushrooms, fungi and insects) (*48*) and lack GH20, GH33 and GH95. The latter GH families, only detected in *B. castoris* and its closest relative *B. italicum*, contain enzymes previously associated with degradation of host-derived carbohydrates in human-associated *Bifidobacterium*: lacto-*N*-biosidases, exo-sialidases and α-L-fucosidases, reported to be involved in metabolism of specific oligosaccharides, including human milk oligosaccharides (HMOs) present in maternal breast milk and intestinal glycoconjugates (*49-52*). Furthermore, the results of the multilevel pattern analysis identified the presence of GH46 chitosanases as the factor driving the differences in GH profiles between strains isolated from *A. agrarius* and the remaining *B. castoris* strains (association index = 0.8819171, *P* < 0.05) (**Supplementary Table S6**). However, only two closely related strains were recovered from this host species, which could explain these results. Overall, these findings highlight a predominance of genes encoding GH families predicted to be responsible for the breakdown of plant-derived polysaccharides in the genomes of *B. castoris* species, and shed light on potential evolutionary adaptations of *B. castoris* strains to the host diet, however with only one non-mouse strain available for the analysis, interpretations can only be tentative.

The glycosyl transferase class of enzymes catalyse the formation of glycosides involved in the biosynthesis of oligosaccharides, polysaccharides and glycoconjugates (*53*) and have previously been associated with production of exopolysaccharide (EPS) in different bacterial species (*54*). A total of 8 GT families were predicted in *B. castoris* genomes, with 18.70±1.61 GT genes on average (21.41±0.02% of predicted glycobiome). GT2 family was predominant in all analysed strains, with an average of 8.15±0.53 GT genes per genome. Carbohydrate esterases, whose function is to release acyl or alkyl groups attached by ester linkage to carbohydrates (*55*), and carbohydrate-binding modules, which have no hydrolytic activity, but bind to carbohydrate ligands and enhance the catalytic efficiency of carbohydrate active enzymes (*55*), constituted 10.66±0.01% and 10.24±0.02% of predicted glycobiome, with 9.33±1.00 and 9.00±1.75 genes per genome, respectively (**Figure 4a and Supplementary Table S5**).

Given the prediction of glycosyl transferase genes in the *B. castoris* glycobiome, we next examined our collection of genomes for the presence of genes potentially involved in exopolysaccharide (EPS) biosynthesis, as previous studies have indicated that EPS may support gut colonisation and stimulate host immune responses (*3*). Recently, relatively conserved genomic regions predicted to contain genes involved in EPS production have been identified in *Bifidobacterium* type strains, including members of *B. pseudolongum* phylogenetic group (gene clusters *eps*3 and *eps*4). For our search, we selected amino acid sequences of *eps* gene clusters from *B. animalis* subsp. *lactis* Bl12 (*eps*3: Bl12_1287 – Bl12_1328) and *B. pseudolongum* subsp. *globosum* LMG 11569^T^ (*eps*4: BPSG_1548 – BPSG_1565) as references (*3*).

This analysis identified homologues of several conserved *eps*-key genes in all analysed *B. castoris* genomes (**Figure 5, Supplementary Table S7**), including those predicted to encode the priming glycosyl transferase (pGTF), which catalyses the first step in EPS biosynthesis, as well as enzymes putatively involved in rhamnose biosynthesis and the transport of the formed EPS-unit across the cytoplasmic membrane (either an ABC-type transporter or a “flippase”-like protein). These results confirm that *B. castoris* strains harbour putative *eps*-key genes and suggest potential ability for this species to produce EPS.

**Figure 5.**
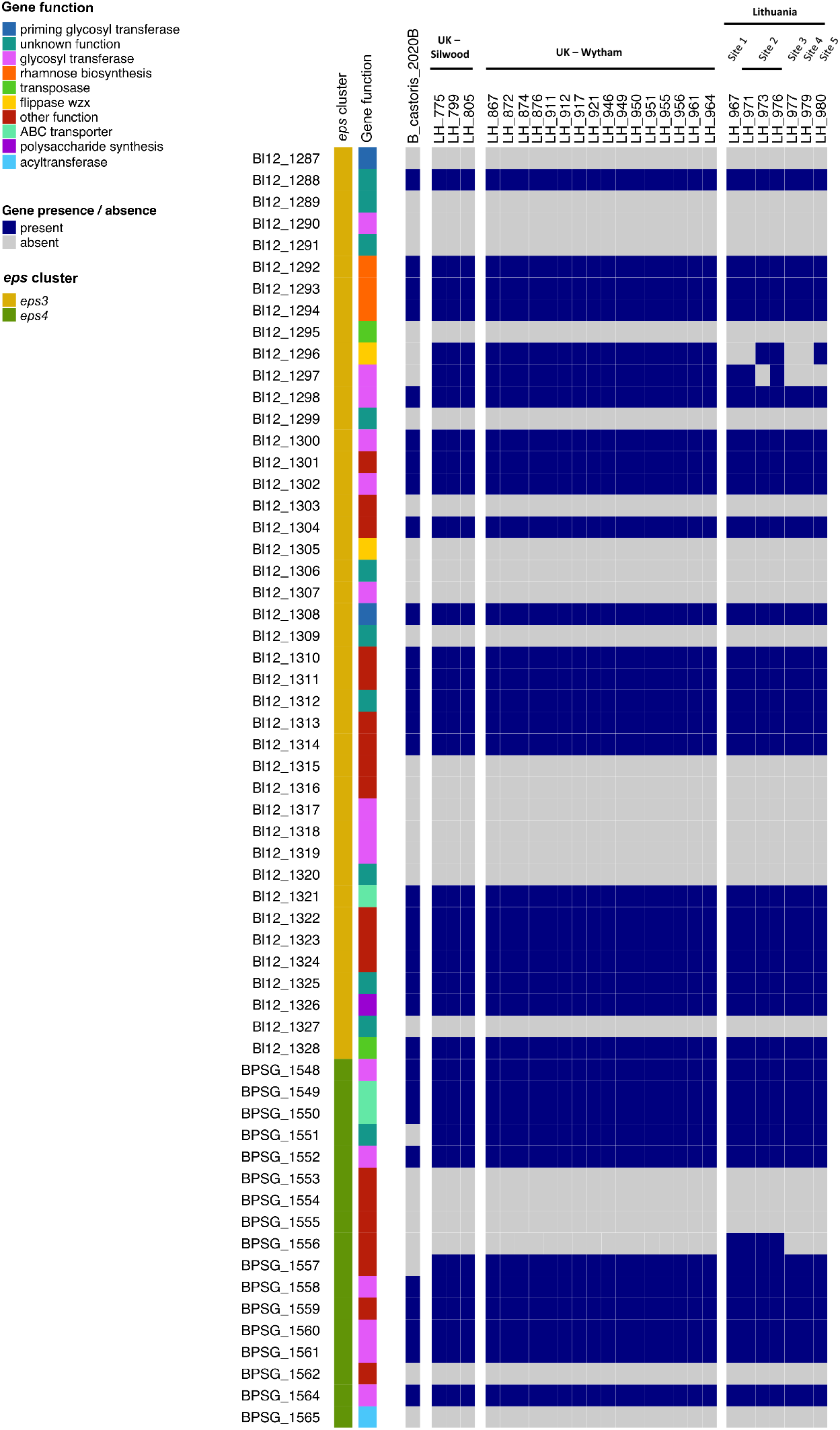
Identification of homologues of *eps*-key genes in *B. castoris*. Sequences of *B. animalis* subsp. *lactis* Bl12 (accession number CP004053.1, *eps*3: Bl12_1287 – Bl12_1328) and *B. pseudolongum* subsp. *globosum* LMG 11569^T^ (accession number JGZG01000015.1, *eps*4: BPSG_1548 – BPSG_1565) were used as reference.

Previously, potential horizontal transfer of *eps* clusters in *Bifidobacterium* has been suggested (*3*). Thus, given the identification of putative *eps*-key genes in our collection of genomes, we next sought to determine the role horizontal gene transfer (HGT) has played in the evolution of *B. castoris* strains. For this purpose, we used the SIGI-HMM tool implemented in software IslandViewer4. This analysis revealed that on average, 6.43±1.15 % of ORFs in *B. castoris* genomes were predicted to be horizontally acquired (range 59 to 191 genes per genome). This value is lower than that previously reported for *B. pseudolongum* species (average of 11.1%), and similar to those reported for *B. animalis* subsp. *lactis* (5.1%) and *B. animalis* subsp. *animalis* (4.6%) (*23, 24*). Cross-referencing with the results of the functional analysis revealed that the eggNOG annotation was available for 49.22±5.69% of putative horizontally acquired genes per genome, on average (**Supplementary Figure 1 and Supplementary Table S8**). The highest proportion of annotated HGT genes in each genome were those of unknown function (38.46±4.36% on average), followed by genes involved in replication, recombination and repair (an average of 19.25±4.43%). This category encompassed CRISPR-Cas-associated proteins, transposases and DNA methylases and methyltransferases. Further analysis of the HGT predictions revealed that genes involved in the cell wall/membrane/envelope biosynthesis constituted on average, 11.35±4.35% of annotated putative HGT genes per genome. This group contained genes neighbouring those identified by our BLAST+ analysis as putative *eps* genes, suggesting they might also be part of *B. castoris eps* clusters (**Supplementary Tables S7 and S8**).

Moreover, genes identified as involved in carbohydrate transport and metabolism constituted on average 3.34±2.10% of predicted annotated horizontally acquired genes per genome. Further analysis of the results for this group revealed that 5 strains from across all locations (the two strains from Silwood Park, one strain from Wytham Woods (LH_950) and two strains from Lithuania (LH_973 and LH_980)) might have acquired a GH43 family member annotated as α-L-arabinofuranosidase through an HGT event. Additionally, strains LH_955 and LH_964 from Wytham Woods, as well as strains LH_973 and LH_979 from Lithuania were predicted to horizontally acquire a GH36 family α-galactosidase (**Supplementary Table S8)**. Overall, these results suggest that HGT may have contributed to the evolution of *B. castoris* strains and their glycobiome, however experimental validation would be essential to confirm the functional importance of these events.

## Discussion

This study is the first to explore strain level genomic signatures of the beneficial bacterial symbiont *Bifidobacterium* in wild rodent populations within an evolutionary and ecological framework. Isolation, whole genome sequencing and in-depth phylogenetic and functional genomic analysis indicates that *B. castoris* appears to be a resident microbiota member of wild rodents belonging to genus *Apodemus*, irrespective of geographical location or host species, with presence of key carbohydrate degradation clusters revealing potential diet-host-microbe evolutionary adaptions.

Our isolation efforts enabled recovery of *B. castoris, B. animalis* and *B. pseudolongum* from three species of wild mouse in the genus *Apodemus* (*A. sylvaticus, A. flavicollis* and *A. agrarius*) from two European countries, UK and Lithuania. Interestingly, despite testing roughly equal numbers of mice and voles captured at the same sites in Lithuania, we did not isolate *Bifidobacterium* from voles. Our inability to detect *Bifidobacterium* in voles could reflect either its absence in these hosts or presence at low abundance, precluding isolation. Although, a recent study that reconstructed metagenome-assembled genomes (MAGs) from faecal metagenomics data from 20 bank voles (*Myodes glareolus*) also indicated an absence of *Bifidobacterium*. Indeed, only one member of the phylum Actinobacteria, among the 254 recovered MAGs, was detected -an organism belonging to the genus *Clavibacter (56)*. However, given the very particular trapping location (Chernobyl Exclusion Zone in Ukraine), it is difficult to speculate on how representative these findings are of vole populations in areas not contaminated with radiation.

Since host specificity represents an avenue of evolutionary development that allows a microbe to colonise a specific host, studying the patterns of host specificity is important for the understanding of the complexities of coevolution between two organisms (*57*). While the widespread distribution of different *Bifidobacterium* species in mammals has previously been postulated, some species seem to show higher host specificity than others. For example, *Bifidobacterium tissieri* and *Bifidobacterium saeculare* have so far only been isolated from primates, and associated with this order based on the analysis of the abundance of bifidobacterial internal transcribed spacer (ITS) rRNA sequences in mammalian faecal samples (*25*). In contrast, *B. animalis* and *B. pseudolongum* had previously been isolated from a number of animal hosts, including mammals and birds, and at the species-level are considered host generalists (*23, 24*). *B. castoris* had previously only been isolated from beavers, but here we show it is also regularly detected in wild mice, which suggests that this species might in fact colonise a broad range of rodents. However additional isolation efforts would be required to confirm its true host range.

Previous research has shown a strong pattern of cophylogeny between some mammalian hosts and *Bifidobacteriaeceae*. Moeller et al. (*11*) showed tight congruence between the phylogenies of *Bifidobacteriaeceae* and their hominid hosts, providing support for codiversification. Similarly, a more recent genome-level cophylogenetic analysis of primate-associated *Bifidobacterium* and their hosts revealed the existence of phylogenetic congruence between the type strains of *Bifidobacterium* typically associated with human hosts and the three members of *Hominidae* family (*Gorilla gorilla, Homo sapiens* and *Pan troglodytes*) (*22*). Here, we test whether a similar pattern holds true based on the analysis of genomic sequences of isolates from a single *Bifidobacterium* species (*B. castoris*) and their wild murine hosts. Moreover, as the species of *Apodemus* are closely related, this provides a different evolutionary perceptive when compared to studies focusing on more distantly related primate species (*35, 58*). In contrast to previous work (*11, 22*), we found no statistical congruence between the *B. castoris* and host phylogenies. *B. castoris* isolates did not show clear phylogenetic clustering by host species or geographic region. Despite this, all *B. castoris* strains identified were only found in a single mouse species, suggesting strains may be host-species specific, but more extensive sampling would be needed to test this definitively. Our data are more consistent with regular host shifts by *B. castoris* strains, and if codiversification has in fact occurred its signature has been eroded by subsequent host shifts.

An interesting question is what might drive the contrasting host-*Bifidobacterium* evolutionary patterns in hominids and mice. We speculate that differences in host evolutionary history and ecology may be important. One possibility is that the level of host-microbe codiversification could differ between members of these two mammalian families. For example, if dietary divergence and associated microbial selection were to be stronger among speciating hominids than *Apodemus*, this could drive stronger codiversification in hominids than mice. Throughout their evolutionary history, the primates have mainly inhabited forest and woodland areas, thus the foods available to them have included the leaves, fruits and flowers of tropical trees and vines (largely dicotyledonous, woody angiosperm species) (*59*). Previous analysis of the dentition of fossil apes (over 15 my old) indicated that they were primarily frugivorous, and to this day herbivorous – and largely fruit-based

– diet dominates among the lesser apes and hominids (*59*). Among the members of *Hominidae*, orangutans and gorillas have been estimated to get around 99% of their annual diet from plant sources, whereas this figure has been lower for chimpanzees, who supplement their ripe fruit diet with selected protein-rich sources that include animal matter – largely invertebrates but occasionally smaller vertebrates (> 87–98%) (*59, 60*). Murine rodents diversified in Europe around 10 mya from the primitive, generalist *Progonomys* that later evolved into lineages related to *Apodemus (61)*. Based on the analysis of the dental pattern of the *Apodemus* mice, it has been suggested that this taxon displayed a relative morphological stability consistent with stabilising selection and retained a primitive, largely granivorous diet over its evolutionary time. These results have been associated with the evolutionary persistence of a forest habitat typically associated with *Apodemus* (mainly deciduous forest that produces seeds), despite the important environmental changes that occurred over the last 10 my (*61*). The more diversified primate diet, compared to that of *Apodemus* species, could perhaps explain the higher *Bifidobacterium*-*Hominidae* specificity.

Alternatively, even if hominids and mice experienced similar codiversification with bifidobacteria initially, variation in the potential for subsequent host shifts may be important. While speciation was likely to have been allopatric for both groups (*11, 62, 63*), allopatry may not have persisted among nascent *Apodemus* species for very long, with higher post-speciation contact among *Apodemus* species allowing more frequent *Bifidobacterium* transfer and host shifts. Hominid species show very strongly bounded present-day geographical ranges (*64*), whereas *Apodemus* species in Europe have very broad overlapping ranges, and different species (e.g. *A. flavicollis* and *A. agrarius* at our Lithuanian sites) can often be caught in adjacent traps. Thus, earlier and more extensive contact between *Apodemus* mouse species may have allowed more cross-species transmission and host shifts of *B. castoris* strains over evolutionary time than could have occurred among hominid species. Future work testing the generality of codiversification across hosts groups with different speciation patterns would be highly informative to understand how the biogeographic and temporal patterns of speciation may affect evolutionary patterns in host-symbiont relationships (*28*).

Characterisation of the *B. castoris* glycobiome provided insight into the strain-specific genetic repertoire predicted to be involved in carbohydrate metabolism and synthesis. The data indicated that *B. castoris* is predominantly enriched in GH families implicated in the degradation of plant-derived carbohydrates. This finding is consistent with the largely plant-based (granivorous) diet of *Apodemus* mice, including as found at several of our sites in UK and Lithuania (*65-68*). The presence of putative family GH49 in *B. castoris* taxon is especially interesting. Most known GH49 dextranases have been discovered in fungi, with only several predicted in bacteria to date (based on CAZy database, July 2020). Overall, members of this family are not very well characterised in prokaryotes (*45*). A recent study using a recombinant dextranase from marine bacterium *Arthrobacter oxydans* KQ11 reported positive effects of its product – an isomalto-oligosaccharide – on the growth of beneficial *Lactobacillus* and *Bifidobacterium*, and the inhibition of pathogenic *E. coli* and *S. aureus* (*45*). However, more studies are required to assess whether the action of enzymes belonging to this GH family could exert beneficial properties on the members on the gut microbiota.

Previously, several bifidobacterial arabinofuranosidases that belong to families GH43, GH51 and GH127 and act on arabinose-substituted polysaccharides have been structurally and functionally identified in *Bifidobacterium* (*69-72*). Our analysis of the GH profiles in *B. castoris* suggested that most strains isolated from mouse hosts lack family GH51 and GH127. Moreover, the data indicate that all *B. castoris* genomes harbour 2-3 copies of GH43 arabinofuranosidases and that some of these copies seemed to have been acquired via an HGT event. These findings suggest there might be an evolutionary advantage for *B. castoris* in possessing GH43 arabinofuranosidases over those belonging to families GH51 and GH127, which may be linked to the composition of the host diet. However, it is difficult to speculate on biological significance of these results without the supporting experimental data. It has previously been shown that arabinofuranosidases characterised in *B. adolescentis* belonging to different families display variation in substrate specificity; AbfA and AXH-d3 belonging to family GH43 removed arabinose on position C(O)2 and C(O)3 of monosubstituted xylose residues and had larger hydrolytic activity towards substrates with a low amount of arabinose substitutions, while AbfB from GH51 only hydrolysed arabinoses from the C(O)3 position of disubstituted xyloses (*71*). The synergistic action of both GH43 arabinofuranosidases has been shown to result in the release of 60% of arabinose from wheat arabinoxylans, which may link to enhanced energy harvest from dietary components associated with low bioavailability (*71*).

Interestingly, three Lithuanian strains isolated from *A. agrarius* and *A. flavicollis* from two distinct trapping sites (site1 and site2) possess genes encoding predicted chitosanases (GH46). The presence of such genes may reflect nutrient availability or dietary preferences of their animal hosts, though no data on the diet of mice analysed in this study were available. Nonetheless, stomach contents analysis has previously detected fungi as a dietary item of *Apodemus flavicollis* in Lithuania at sites close to those studied here (*65*) as well as in one of our UK sampling sites (*66*), and spores of putatively edible fungi (with macroscopic fruiting bodies) were frequently detected in the faeces *Apodemus* spp. in other parts of Lithuania (*73*), suggesting mycophagy in *Apodemus* spp. may not be uncommon. Further information on the specific food items eaten by *Apodemus* could allow bioinformatic predictions of carbohydrate degradation properties of *B. castoris* strains, based on specific dietary components found in host diet. The potential of *Bifidobacterium* to degrade chitin-derived substrates is currently not very well understood. Previously, studies looking into functional effects of chito-oligosaccharides (COS) on gut microbiota members produced inconsistent results. Lee et al. (*74*) showed increased growth of *B. bifidum* in pure cultures supplemented with COS. Contrary to this, Vernazza et al. (*75*) did not observe any positive effects of COS on growth of human-associated bifidobacteria from faecal inocula. Furthermore, Yang et al. (*76*) reported an increase in the population of *Bifidobacterium* upon dietary supplementation of weaning pigs with COS, however a recent study using both in *vitro* fermentation and *in vivo* mouse model indicated that *Bifidobacterium* growth was significantly inhibited in mice fed with COS, leading to the conclusion that these oligosaccharides should not be considered preferred prebiotic substrates (*77*).

These same Lithuanian strains, unlike all the remaining *B. castoris* strains, also appear to be lacking GH families containing enzymes previously associated with degradation of specific fucosylated and sialylated milk oligosaccharides and intestinal glycoconjugates in *Bifidobacterium* isolated from human hosts (GH20, GH33, GH95). The structure and composition of the milk oligosaccharides differ greatly between mammalian species. According to Prieto et al. (*78*) human milk contains the most complex mixture of reducing oligosaccharides, many of which include fucose and determinants for human blood groups, e.g. the ABO and Lewis system. Compared to human breast milk, composition of oligosaccharides in mouse milk is mainly limited to sialyllactoses with minuscule amounts of fucosylated lactose (3’-fucosyllactose) (*78*). The fact that we identified enzymes belonging to families GH20, GH33 and GH95 in *B. castoris* strains indicate that members of this species may have the ability to metabolise specific host-derived oligosaccharides, including milk oligosaccharides, however experimental evaluation of substrate specificity of these enzymes is required.

The identification of genes predicted to encode glycosyl transferases (GTs) in genomes of *B. castoris* prompted questions about potential ability of members of this species to produce EPS. This bifidobacteria trait is associated with immune modulation and longer-term persistence in the (laboratory) murine gastrointestinal tract, suggesting EPS may also play a key role in microbe-host interactions in wild mice populations (*79-81*). Recent genomic studies on *Bifidobacterium* type strains have described high levels of inter-species variation with respect to the number, function and organisation of genes in *Bifidobacterium eps* clusters (*3, 54*). However, a set of conserved *eps*-key genes has been proposed as universal markers, including genes predicted to encode the priming glycosyl transferase (pGTF), other glycosyl transferases, transporter enzymes (either “flippases” or ABC transporters) and various carbohydrate precursor biosynthesis or modification enzymes (*54, 82*). The results of our BLAST+ search for the homologues of these *eps*-key genes previously identified in *Bifidobacterium* type strains most closely related to *B. castoris* 2020B^T^ revealed their presence in *B. castoris* isolates, suggesting this species may be able to synthesize EPS. Furthermore, the analysis of predicted HGT events in our isolates identified additional genes of unknown function neighbouring the *eps*-key genes that may be part of a distinct *B. castoris eps* cluster. These findings support previous suggestions on possible role of HGT in acquisition of complete or partial *eps* clusters by *Bifidobacterium* (*3*). In line with these observations, we identified homologues of protein members of clusters *eps3* and *eps4* in *B. castoris*, including enzymes involved in rhamnose biosynthesis in cluster eps3 which may link to additional biological properties of these polymers (*83, 84*). However, additional studies are required to assess the functionality of the putative EPS biosynthesis machinery in *B. castoris* species.

## Conclusion

It is well recognised that members of *Bifidobacterium* exert beneficial health effects on their host, however current knowledge of their diversity, distribution across the host phylogeny, and metabolic capability in non-human hosts, especially in wild animal populations, is limited. This research provides novel insights into the host-microbe evolutionary relationships and genomic features of *B. castoris* isolated from geographically distinct wild mouse populations. Our initial observations on strain-specific carbohydrate metabolism repertoires and the presence of *eps* genes require further investigation to understand how *Bifidobacterium* adapts, persists and interacts with the animal host and explain the functionality of mechanisms underlying bifidobacterial metabolic activity. This could be achieved through a combination of experimental and *in silico* methods, including additional isolation experiments and whole genome sequencing, combined with community analyses based on metagenomic approaches, as well as carbohydrate metabolism and transcriptomics assays.

## Materials and methods

### Faecal sample collection

In the UK, live rodent trapping was carried out at two sites of mixed deciduous woodland approximately 50km apart: Wytham Woods (51° 46’N,1°20’W) and Nash’s Copse, Silwood Park (51°24’N, 0°38’W). All animals were live-trapped using a standard protocol across both sites, using small Sherman traps baited with peanuts and apple and provisioned with bedding, set at dusk and collected at dawn the following day. All newly captured individuals were marked with subcutaneous PIT-tags for permanent identification, and all captures were weighed and various morphometric measurements taken. Faecal samples were collected using sterilised tweezers from the base of Sherman traps into sterile tubes. Samples from Silwood were frozen within 8 hours of collection at -80°C and sent frozen to the Quadram Institute (Norwich, UK) for *Bifidobacterium* culturing; samples from Wytham were posted on the day of sampling at room temperature to the same address for culturing. Upon reception of samples, they were immediately frozen at -80°C. To ensure no cross-contamination and identification of samples to specific individuals, any traps that showed signs of rodent presence (captures and trigger failures) were washed thoroughly in a bleach solution and autoclaved between uses.

In Lithuania, small mammals were trapped in October 2017 and between May -November in 2018 using live-and snap-traps at twelve locations: Site 1: 54.92878, 25.33333; Site 2: 55.02814, 25.27380; Site 3: 54.92866, 25.31524, Site 4: 55.05938, 25.35643, Site 5: 54.96362, 25.35640; Site 6: 54.93026, 25.24138; Site 7: 54.99276, 25.24909, Site 8: 55.02700, 25.35867; Site 9: 55.06756, 25.29782; Site 10: 54.76482, 25.31283; Site 11: 54.76322, 25.35052; Site 12: 54.93650, 25.28442. Traps baited with bread soaked in sunflower oil (in case of live-traps, apple and bedding were also added) were set in the evening and retrieved in the morning. Small mammals trapped with snap-traps were placed in separate bags and transported to the lab on ice. Live-trapped animals were transported to the lab and humanely killed by cervical dislocation. Species, sex, age, reproduction status of small mammals were identified. Content of distal part of colon (∼20-30 mm) was removed, placed in Eppendorf tube and stored at –80°C. Frozen samples were sent to the Quadram Institute (Norwich, UK) for *Bifidobacterium* culturing.

The distance between trapping sites in both the UK and Lithuania was far enough for the animals not to move between them. All studied species have small home ranges – *Apodemus* spp. have the widest range and rarely move more than 0.25 km (*85*).

### Strain isolation

Depending on the number of available faecal pellets, samples were re-suspended in either 450ul or 900ul of sterile Phosphate Buffer Saline (PBS) (Sigma-Aldrich, UK) and used to produce 10-fold serial dilutions (neat -10^−4^). The samples were then vortexed for 30s and mixed using on a shaker at 1600rpm. An aliquot of each dilution (100 ul) was plated onto Brain Heart Infusion (BHI) (Oxoid, UK) agar supplemented with mupirocin (50mg/l) (Sigma-Aldrich, UK), L-cysteine hydrochloride monohydrate (50mg/l) (Sigma-Aldrich, UK) and sodium iodoacetate (7.5mg/l) (Sigma-Aldrich, UK) and incubated in an anaerobic cabinet for 48-72 hours. Three colonies from each dilution were randomly selected and streaked to purity on BHI agar supplemented with L-cysteine hydrochloride monohydrate (50mg/l). Pure cultures were stored in cryogenic tubes at -80°C.

### DNA extraction, whole-genome sequencing, assembly and annotation

DNA for whole genome sequencing was extracted from pure bacterial cultures using the phenol-chloroform method as described previously (*36*), and subjected to multiplex Illumina library preparation protocol and sequencing on Illumina HiSeq 2500 platform at the Wellcome Trust Sanger Institute (Hinxton, UK) or Illumina MiSeq at the Quadram Institute Bioscience (Norwich, UK) (**Supplementary Table S1**). Sequencing reads were screened for contamination using Kraken v1.1 (MiniKraken) (*86*) and pre-processed with fastp v0.20 (*87*). SPAdes v3.11 with “--careful” option (*88*) was used to produce assemblies, after which contigs below 500bp were filtered out. Additionally, publicly available genome assembly of *Bifidobacterium* castoris 2020B^T^ type strain (accession number NZ_QXGI00000000.1) was retrieved from NCBI Genome database (*4*) and all genomes were annotated with Prokka v1.13 (*89*). The draft genomes of 50 *Bifidobacterium* isolates sequenced here have been deposited to GOLD database at https://img.jgi.doe.gov, GOLD Study ID: Gs0153956.

### Pangenomic and phylogenomic analysis

Anvi’o version 6.1 (*90*) was used to generate *B. castoris* pangenome and single copy core gene data for other analyses. Briefly, we created a text file containing required information on our collection of genomes and used this file to generate genomes storage database. We then computed the pangenome using the script anvi-pan-genome with parameters “--minbit 0.5 --mcl-inflation 10 --use-ncbi-blast”. We next identified the single copy core genes in the pangenome and recovered their aligned amino acid sequences using with anvi-get-sequences-for-gene-clusters with parameters “--min-num-genomes-gene-cluster-occurs, --max-num-genes-from-each-genome, --concatenate-gene-clusters”. The resulting output was cleaned from poorly aligned positions using trimAl v1.4.1 (gaps in more than 50% of the genes) (*91*). IQ-TREE v1.6.1 (*92*) employing the ‘WAG’ general matrix model with 1000 bootstrap iterations (*93*) was used to infer the maximum likelihood trees from protein sequences. The host tree for cophylogentic analysis was constructed using concatenated sequences for 12S rRNA and partial ctyb genes employing the ‘GTR’ model with 1000 bootstrap iterations. ParaFit (*34*) in the ‘ape’ package of R (*34, 94*) was used for distance-based comparisons, with the code included in the Supplementary Table S9. Python3 module pyANI v0.2.10 with default BLASTN+ settings was used to calculate the average nucleotide identity (ANI) (*95*). Species delineation cut-off was set at 95% identity (*96*). Isolates showing identity values above 99.9% were considered identical (*29*). We used the script anvi-import-misc-data to import the results the anvi’o pangenome database and visualised the output with anvi-display-pan.

### Comparative genomics

Functional categories (COG categories) were assigned to genes using EggNOG-mapper v0.99.3, based on the EggNOG database (bacteria) (*97*) and the abundance of genes involved in carbohydrate metabolism was calculated. Standalone version of dbCAN2 (v2.0.1) was used for glycobiome prediction (*98*). ‘Vegan’ and ‘indicspecies’ packages implemented in R were used for the plotting of the *B. castoris* GH profiles using non-metric multidimensional scaling and for the multilevel pattern analysis with the point biserial correlation coefficient, respectively (*99, 100*), with the code included in the Supplementary Table S9. BLAST+ v2.9.0 (e-value of 1e-5 and 50% identity over 50% sequence coverage) (*101*) was used to screen *B. castoris* genomes for the presence of homologues of *eps* genes from *B. animalis* subsp. *lactis* Bl12 (accession number CP004053.1, *eps*3: Bl12_1287 – Bl12_1328) and *B. pseudolongum* subsp. *globosum* LMG 11569^T^ (accession number JGZG01000015.1, *eps*4: BPSG_1548 – BPSG_1565). SIGI-HMM (*102*) tool implemented in Islandviewer4 (*103*) was employed to predict HGT events.

## Acknowledgments

The authors would like to thank Dr Florent Mazel for insightful discussions and providing feedback on the study. This work was funded by a Wellcome Trust Investigator Award (no. 100/974/C/13/Z); a BBSRC Norwich Research Park Bioscience Doctoral Training grant no. BB/M011216/1 (supervisor LJH, student MK); an Institute Strategic Programme Gut Microbes and Health grant no. BB/R012490/1 and its constituent projects BBS/E/F/000PR10353 and BBS/E/F/000PR10356; and an Institute Strategic Programme Gut Health and Food Safety grant no. BB/J004529/1 to LJH. SCLK was funded by a NERC fellowship (NE/L011867/1), and AR by a Clarendon Scholarship from the University of Oxford. LB was funded by grant S-MIP-17-86 from the Research Council of Lithuania. The funding bodies did not contribute to the design of the study, collection, analysis, and interpretation of data or in writing the manuscript.

## Author contributions

LJH and MK designed the overall study. AR, LB and SCLK provided the faecal samples from wild mice populations and associated metadata. MK isolated the strains, extracted the DNA and prepared it for WGS, performed all genomic analysis and visualisation. MK and LJH analysed the data, with input and discussion from SCLK and AR, and drafted the manuscript. AR, LB and SCLK provided further edits and co-wrote the final version. All authors read and approved the final manuscript.

---

Global test: ParaFitGlobal = 8.835461e-05, p-value = 0.3586 (9999 permutations) Test of individual host-parasite links (9999 permutations)

|  | Host | *B. castoris* isolate | F1.stat | p.F1 | F2.stat | p.F2 |
| --- | --- | --- | --- | --- | --- | --- |
| [26,] | A. agrarius | LH_967 | 2.66E-05 | 0.3073 | 9.07E-03 | 0.3068 |
| [27,] | A. agrarius | LH_971 | 2.66E-05 | 0.3064 | 9.08E-03 | 0.3064 |
| [21,] | A. flavicollis | LH_980 | -6.22E-07 | 0.7765 | -2.12E-04 | 0.7765 |
| [22,] | A. flavicollis | LH_976 | -1.76E-05 | 0.808 | -5.99E-03 | 0.8093 |
| [23,] | A. flavicollis | LH_973 | -1.52E-05 | 0.7834 | -5.17E-03 | 0.7849 |
| [24,] | A. flavicollis | LH_977 | -2.70E-08 | 0.5686 | -9.20E-06 | 0.5686 |
| [25,] | A. flavicollis | LH_979 | 7.89E-09 | 0.5594 | 2.69E-06 | 0.5594 |
| [2,] | A. sylvaticus | LH_775 | 1.14E-06 | 0.4388 | 3.90E-04 | 0.4365 |
| [3,] | A. sylvaticus | LH_799 | 1.14E-06 | 0.4455 | 3.90E-04 | 0.4446 |
| [4,] | A. sylvaticus | LH_950 | -3.04E-06 | 0.8379 | -1.04E-03 | 0.8376 |
| [5,] | A. sylvaticus | LH_805 | 6.00E-06 | 0.2779 | 2.04E-03 | 0.2764 |
| [6,] | A. sylvaticus | LH_867 | 6.98E-06 | 0.2134 | 2.38E-03 | 0.2133 |
| [7,] | A. sylvaticus | LH_872 | 6.99E-06 | 0.2109 | 2.38E-03 | 0.2114 |
| [8,] | A. sylvaticus | LH_876 | 6.98E-06 | 0.2105 | 2.38E-03 | 0.211 |
| [9,] | A. sylvaticus | LH_911 | 6.99E-06 | 0.2112 | 2.38E-03 | 0.2117 |
| [10,] | A. sylvaticus | LH_946 | 6.99E-06 | 0.212 | 2.38E-03 | 0.2125 |
| [11,] | A. sylvaticus | LH_949 | 6.99E-06 | 0.2096 | 2.38E-03 | 0.2097 |
| [12,] | A. sylvaticus | LH_961 | 6.99E-06 | 0.2187 | 2.38E-03 | 0.2189 |
| [13,] | A. sylvaticus | LH_956 | 6.99E-06 | 0.2164 | 2.38E-03 | 0.2166 |
| [14,] | A. sylvaticus | LH_951 | 6.99E-06 | 0.2112 | 2.38E-03 | 0.2116 |
| [15,] | A. sylvaticus | LH_874 | 7.00E-06 | 0.2105 | 2.39E-03 | 0.2102 |
| [16,] | A. sylvaticus | LH_912 | 6.99E-06 | 0.2121 | 2.38E-03 | 0.2131 |
| [17,] | A. sylvaticus | LH_917 | 7.21E-06 | 0.2088 | 2.46E-03 | 0.2093 |
| [18,] | A. sylvaticus | LH_921 | 7.21E-06 | 0.214 | 2.46E-03 | 0.2135 |
| [19,] | A. sylvaticus | LH_955 | 3.41E-06 | 0.4128 | 1.16E-03 | 0.4104 |
| [20,] | A. sylvaticus | LH_964 | 3.41E-06 | 0.4074 | 1.16E-03 | 0.4049 |
| [1,] | B. castoris | B_castoris_2020B | 2.75E-06 | 0.3883 | 9.36E-04 | 0.3854 |

