## Supplementary Figure 1 for "*Bifidobacterium castoris* strains isolated from wild mice show evidence of frequent host switching and diverse carbohydrate metabolism potential"

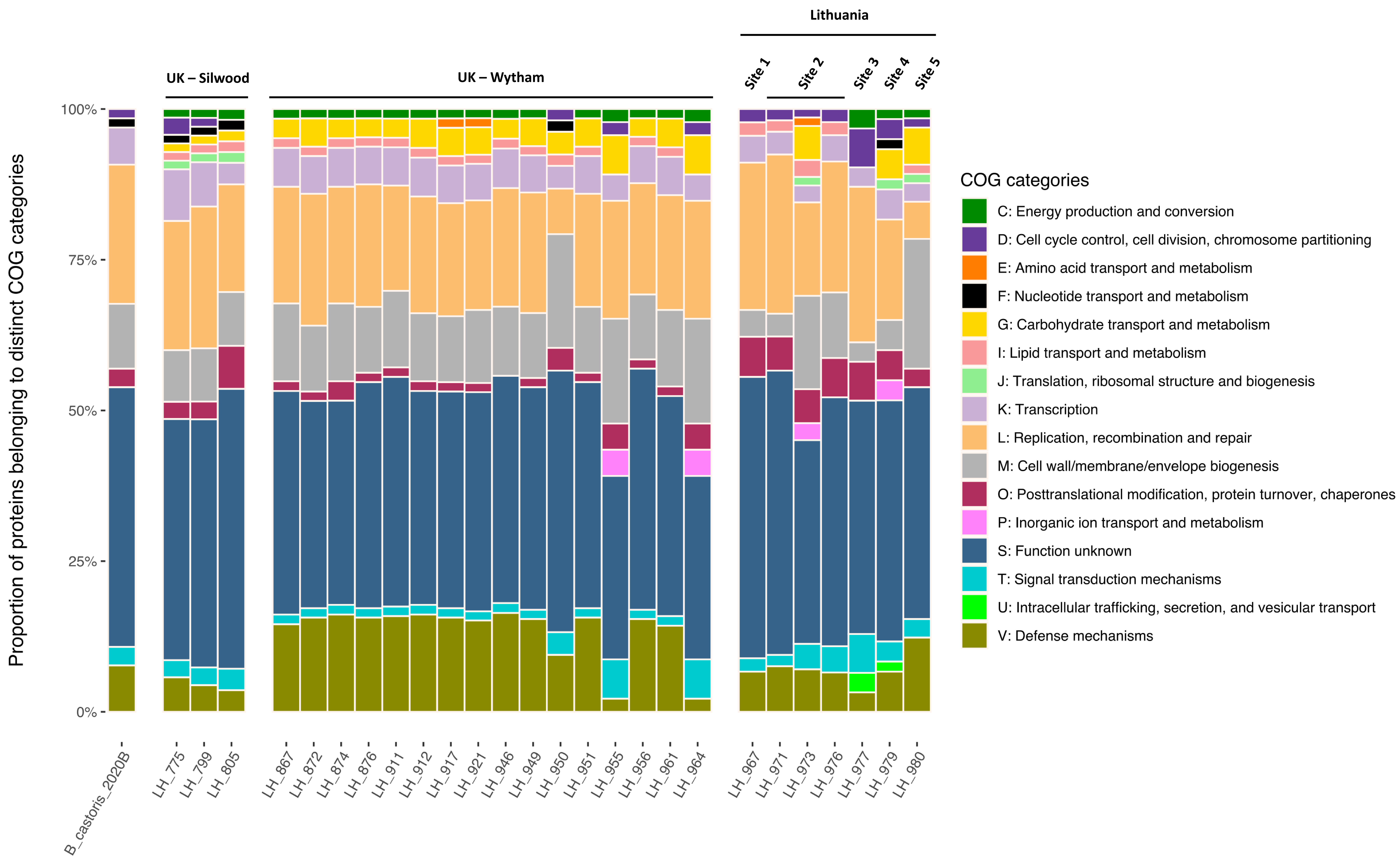


Supplementary Figure 1. Functional classification of proteins predicted to be horizontally acquired by *B. castoris* isolates according to COG categories based on the available eggNOG annotation. The eggNOG annotation was available for 49.22±5.69% of putative horizontally acquired genes per genome, on average.
